# From exact gradients to exact standard errors: third- and fourth-order sensitivities for the Laplacian population likelihood family

**DOI:** 10.64898/2026.07.09.737434

**Authors:** H. van de Beek, M. Beldjenna, J. Almquist, M. Fidler, L.B. Zwep, J.G.C. van Hasselt

## Abstract

Asymptotic standard errors for the parameters of a nonlinear mixed-effects model fitted by Laplacian estimation, first-order conditional estimation (FOCE), or FOCE with interaction (FOCEI) require the observed (Fisher) information, the negative second derivative of the population objective at the optimum. The gradient of this objective can be computed exactly from sensitivity equations, but the observed information is still formed by finite differencing, which is less accurate, step-size dependent, and requires empirical Bayes reoptimization. These perturbed individual optimization problems can fail even after a successful fit. Our objectives are to extend the exact-gradient sensitivity framework by one differentiation and give the observed information of the Laplacian population likelihood family in closed form and (2) illustrate covariance calculation without further empirical Bayes optimization in a case example. Writing the objective as a data term plus the log-determinant of the inner Hessian, the population Hessian splits so that the data term reuses second-order sensitivities, whereas the log-determinant term requires third-order sensitivities for FOCE/FOCEI and fourth-order sensitivities for the full Laplacian method. We illustrate this on a two-compartment oral model with first-order absorption fitted to simulated data. The analytic covariance involves no differencing step and uses the fitted empirical Bayes estimates without reoptimization, avoiding this source of covariance-step failure. Implemented for FOCE/FOCEI in the open-source R package nlmixr2, the construction provides exact, reproducible standard errors for the approximate likelihood, limited by numerical solution accuracy and conditioning. This supports more dependable confidence intervals, identifiability assessment, and uncertainty propagation.

## Introduction

Nonlinear mixed-effects (NLME) models are the standard framework for population pharmacokinetic and pharmacodynamic analysis, in which observations are pooled across individuals through a population distribution of random effects. Estimation by the full Laplacian method, first-order conditional estimation (FOCE), and FOCE with interaction (FOCEI) maximizes an approximation to the population likelihood [1, 2]. Inference about the estimated parameters — confidence intervals and identifiability diagnostics — rests on their asymptotic standard errors, which are obtained from the observed (Fisher) information, the negative second derivative of the population objective with respect to the parameters, evaluated at the optimum [3].

Both the gradient and the Hessian of this objective have traditionally been formed by forward or central finite differencing [4], whose noise and step-size sensitivity can be problematic. The gradient can now be computed exactly from first- and second-order sensitivity equations for FOCE/FOCEI, and additionally third-order sensitivities for the full Laplacian method [5], removing that source of error from the point estimates and from the optimization that produces them. The corresponding observed information, however, has not been given in closed form, and is conventionally still obtained by finite differencing [6, 7]. The second derivative, for which differencing is markedly less accurate than for a gradient, can therefore be even more problematic. Each perturbed objective evaluation also requires solving the empirical Bayes problems again, so a successful fit can be followed by a failed covariance step if a perturbed individual optimization fails.

The differentiation order is governed by the inner Hessian, which under FOCE/FOCEI is the first-order approximation built from first-order sensitivities: differentiating the objective once introduces second-order sensitivities (the gradient), and differentiating it twice introduces third-order sensitivities. The full Laplacian method uses the exact inner Hessian, built from first- and second-order sensitivities, and therefore requires third-order sensitivities for the gradient and fourth-order sensitivities for the observed information. Written as a data term plus a log-determinant term, the observed information splits so that the higher-order dependence is confined to the log-determinant term; the data term requires nothing beyond the second-order sensitivities already needed for the gradient.

In this study we aim to (1) extend the exact-gradient sensitivity framework by one differentiation and give the observed information of the Laplacian population likelihood family in closed form and (2)illustrate covariance calculation without further empirical Bayes optimization in a case example.

### Theory

#### Model and objective

Let subjects *i* (*i* = 1, …, *N*) be described by a structural model

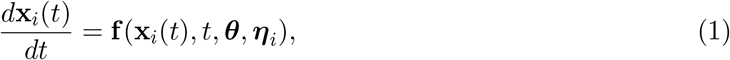

with initial state

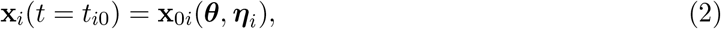

where **x**_*i*_ is the vector of state variables for the i-th individual, ***θ*** is a vector encompassing fixed effect parameters, residual-error parameters and covariances of random effect parameters (with covariance matrix **Ω**), and ***η***_*i*_ ∼ *N* (0, **Ω**) is the vector of random effects for the i-th individual.

For *j* (*j* = 1, …, *n*_*i*_), **y**_*ij*_ describes the j-th observation for the i-th individual and follows the observation model

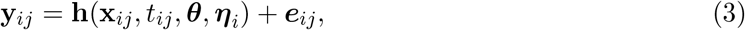

where ***e***_*ij*_ ∼ *N* (0, **R**_*ij*_(**x**_*ij*_, *t*_*ij*_, ***θ, η***_*i*_)) is residual unexplained variability, and **h**(**x**_*ij*_, *t*_*ij*_, ***θ, η***_*i*_) = *E*(**y**_*ij*_|***η***_*i*_) = **ŷ**_*ij*_ is the individual prediction.

Given experimental observations **d**_*ij*_ (realization from **y**_*ij*_) and ***ϵ***_*ij*_ = **d**_*ij*_ − **ŷ**_*ij*_ the associated residuals, the population likelihood can be expressed as

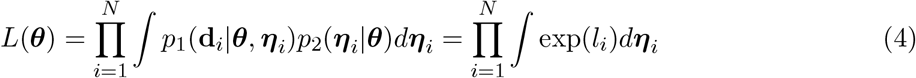

where *l*_*i*_ is the individual joint log-likelihood of the data encompassing both the data term and the random effect term

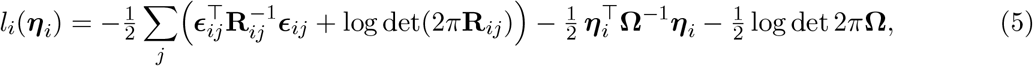

The population likelihood (4) does not have an exact closed form solution, but it can be approached by approximating *l*_*i*_ with a second order Taylor expansion with respect to ***η***_*i*_ around the empirical Bayes estimate 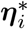 that maximises 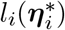. This is the Laplacian approximation.

With 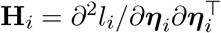 the exact inner Hessian, the population objective is the Laplacian approximation

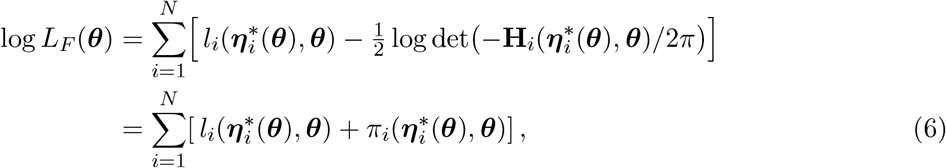

with for each subject the data term 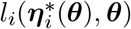 and the penalty 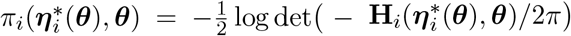.

Asymptotic standard errors follow from the observed information − *d*^2^ log *L*_*F*_ */d****θ*** *d****θ***^⊤^ at the optimum, with 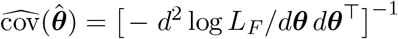 (optionally the sandwich form with the cross-product matrix).

Throughout we assume that **f**, **x**_0*i*_, **h, R**_*ij*_ and **Ω** are four times continuously differentiable in (***θ, η***_*i*_), that **R**_*ij*_ *>* 0 and **Ω** ≻ 0, and that for each ***θ*** the inner problem has a unique strict local maximizer 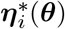 at which the exact inner Hessian **H**_*i*_ is nonsingular on a neighbourhood. The implicit function theorem then makes 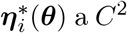 map, and the smoothness of the forward sensitivity solutions allows us to interchange element-wise partial derivatives in the ***θ***-derivations below.

The gradient of this Laplacian objective is

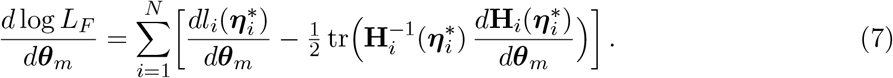

For FOCE/FOCEI, the same form holds with 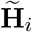 and its derivative replacing **H**_*i*_ and *d***H**_*i*_*/d****θ***_*m*_ in the trace term. The exact inner Hessian is retained when differentiating the individual optimality condition below. Since 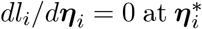, the first term reduces to the explicit partial *∂l*_*i*_*/∂****θ***_*m*_. The same simplification does not apply to the second term, for which we require the indirect dependence through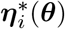. Differentiating the optimality condition gives the mode derivative,

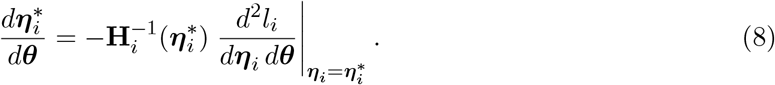

We can then obtain the observed information by differentiating once more (7) with respect to ***θ***; the remainder of this section evaluates it in closed form.

#### The observed information

The second total derivative of the log likelihood with respect to ***θ*** — through 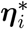 as well as explicitly — gives the population Hessian

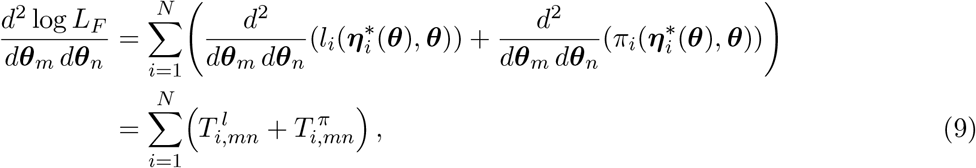

The labels *T* ^*l*^, *T* ^*π*^ name only the second derivatives of the two summands of (6) and bear no relation to the method names (FOCE, FOCEI, Laplacian).

Both terms use the mode derivatives. Equation (8) gives the first, and differentiating once more gives the second (derivation in Appendix):

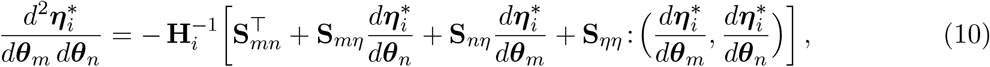

where **S**_· ·_ = *d*^3^*l*_*i*_*/d*(·) *d*(·) *d****η***_*i*_ are the third derivatives of *l*_*i*_, the trailing differentiation always in ***η***_*i*_ (from *dl*_*i*_*/d****η***_*i*_ = 0) and the two subscripts naming the other directions: **S**_*mn*_, where *m, n* designate directions ***θ***_*m*_, ***θ***_*n*_, is a row vector in ***η*, S**_*mη*_, where *η* designates direction ***η***_*i*_, is a matrix, and **S**_*ηη*_ a rank-three tensor symmetric in its two ***η*** slots. The colon denotes the bilinear application (*A* :(*u, v*))_*µ*_ = Σ_*k,l*_ *A*_*µkl*_ *u*_*k*_*v*_*l*_.

**Data term**. Since *dl*_*i*_*/d****η***_*i*_ = 0 at 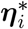, differentiating the data term *l*_*i*_ twice gives (derivation in Appendix):

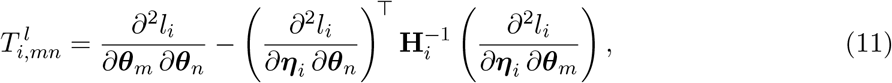

which introduces no new sensitivity order: the mixed 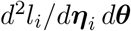 is reused from the gradient (7), and the pure 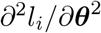 is a second-order quantity, formed here for the first time but requiring no sensitivity beyond second order.

#### Log-determinant term

With *π*_*i*_ the penalty as defined in (6), differentiating it twice gives (derivation in Appendix)

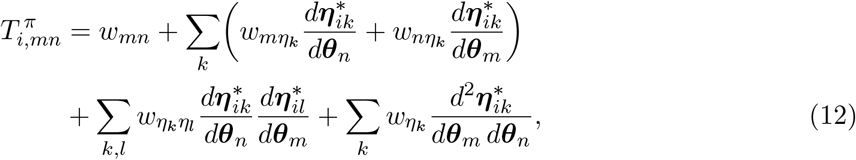

with the primitives, for *s, t* ∈ {*m, n, η*_*k*_, *η*_*l*_} denoting the directions {***θ***_*m*_, ***θ***_*n*_, ***η***_*ik*_, ***η***_*il*_}, respectively,

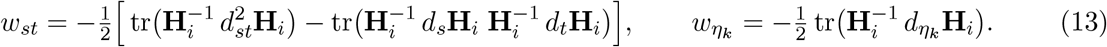

### Reparameterization

Equations (9)–(13) are written for the parameter vector ***θ***. When estimation is carried out in an unconstrained reparameterization ***θ*** = ***θ***(***β***) (for example a log scale for positive parameters, or a Cholesky factor for **Ω**), with *a, b* indexing the components of ***β***, the Hessian transforms by the chain rule,

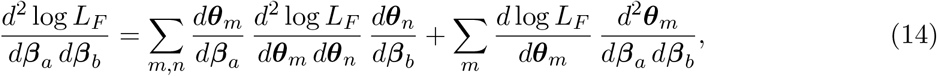

the second sum vanishing for affine maps and equal to the gradient times *d*^2^***θ***_*m*_*/d****β***^2^ otherwise.

### FOCE and FOCEI

FOCE and FOCEI are based on this Laplacian approximation, where the inner Hessian **H**_*i*_ = Δ*l*_*i*_ is further simplified by approximating various terms by their expected values over the data, conditional on ***η***_*i*_. There, the observations are considered as random variables whose values have not yet been realized which gives *E*[***ϵ***_*ij*_ | ***η***_*i*_] = 0 and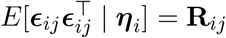.

The NONMEM variant is written

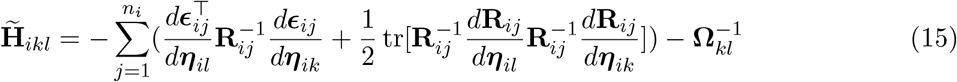

Since *ϵ*_*ij*_ = d_*ij*_ − ŷ _*ij*_,

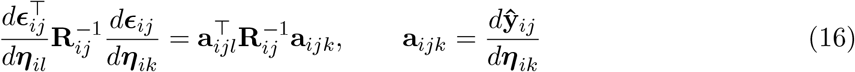

Using the invariability of the trace under cyclic permutation, and since 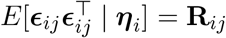, we can rewrite (derivation in Appendix):

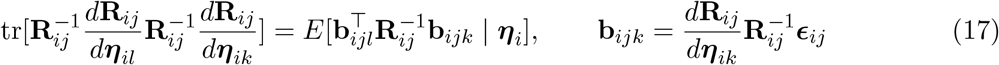

The expectation is over data at fixed ***θ*** and ***η***_*i*_, and is taken before evaluating at the mode. Hence the following expression of the FOCEI inner Hessian

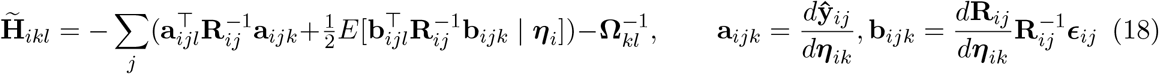

in which terms are approximated to their conditional expected values while the full Laplacian method does not use this approximation. Here **a**_*ijk*_ are the first-order prediction sensitivities, while **b**_*ijk*_ is the interaction contribution and is dropped under FOCE with residual covariance independent of ***η*** . 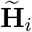 is negative definite because the quadratic contributions are positive semidefinite and **Ω**^−1^ is positive definite.

For FOCE/FOCEI, **H**_*i*_ replaces 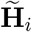 in the log-determinant of (6), the trace term of (7), and the primitives (13). The exact inner Hessian is retained in the mode derivatives (8) and (10) and in the data term (11). The derivation above otherwise applies unchanged.

Under the FOCE approximation, the differentiation order can be exposed by writing the derivatives of (18) in the sensitivities:

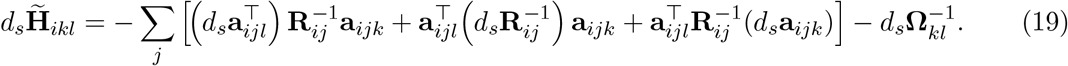

with *d*_*s*_**a**_*ijk*_ = *d*^2^**ŷ**_*ij*_*/ds d****η***_*ik*_ and 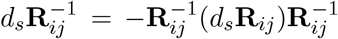. One more differentiation of (19) makes 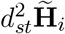 carry 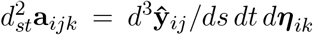, third order for directions *s, t*, ***η***_*ik*_, as well as the second derivatives of **R**_*ij*_ (under FOCEI, the corresponding third derivatives of **R**_*ij*_ additionally appear through the trace term in (15)). The third order therefore enters in exactly two places, both in *T π*: the explicit 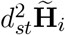of (13), and the second mode derivative (10) in the last term of (12).

#### Order of the sensitivity equations

Under the full Laplacian approximation, **H**_*i*_ already carries *d*^2^**ŷ***/d****η***^2^. The gradient differentiates it once, requiring third-order sensitivities; the observed information differentiates it twice, requiring fourth-order sensitivities *d*^4^**ŷ***/d****η***^4^, *d*^4^**ŷ***/d****η***^3^*d****θ***, and *d*^4^**ŷ***/d****η***^2^*d****θ***^2^, with the corresponding residual-covariance derivatives. The fourth order enters only in *d*^2^ **H**_*i*_ of (13); the second mode derivative (10) remains third order in *l*_*i*_.

For FOCE/FOCEI, the differentiation order is set by 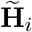 of (18), built from the first-order sensitivities *d***ŷ**_*ij*_*/d****η***_*ik*_ . The gradient (7) differentiates 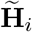 once, introducing the second-order sensitivities *d*^2^**ŷ***/d****η***^2^ and *d* **ŷ***/d****η*** *d****θ***. The observed information differentiates 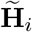 a second time, raising each sensitivity by one order: it requires the third-order sensitivities *d*^3^**ŷ***/d****η***^3^, *d*^3^**ŷ***/d****η***^2^*d****θ***, and *d*^3^**ŷ***/d****η*** *d****θ***^2^ (with the corresponding third derivatives of **R**_*ij*_ under FOCEI), that is, third-order sensitivity equations. The third-order sensitivities used here are obtained exactly by extending the sensitivity equations to third order, which adds one chain-rule term per elementary operation,

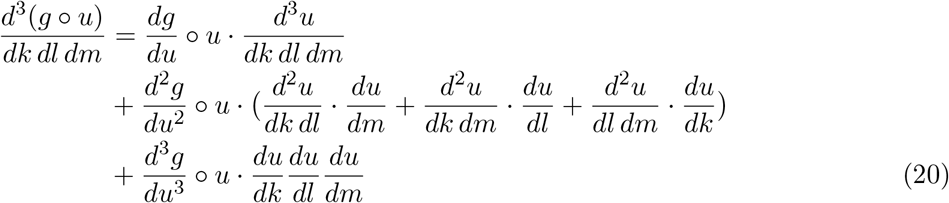

The fourth-order chain rule, adding direction *n* to the same *k, l, m*, is

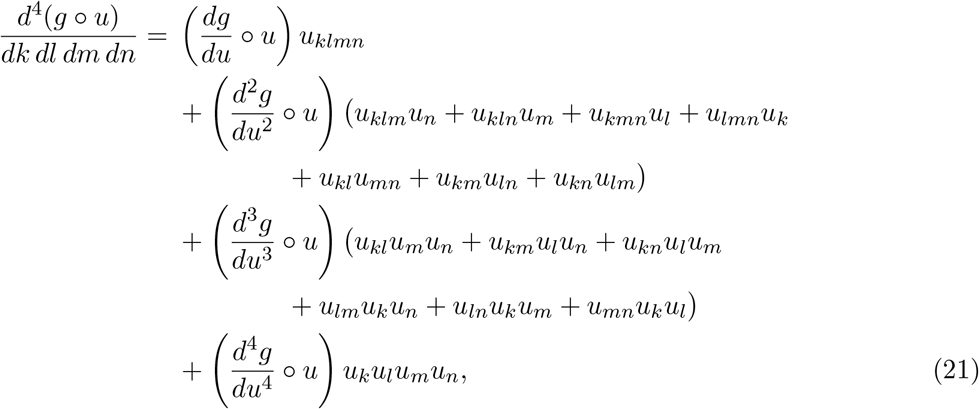

where *u*_*k*_ = *du/dk, u*_*kl*_ = *d*^2^*u/dk dl*, and similarly for higher orders. Thus (20) supplies the third order needed for FOCE/FOCEI observed information and the full Laplacian gradient; (21) supplies the fourth order additionally needed for full Laplacian observed information.

This set reduces under mu-referencing. When a structural parameter ***θ***_*k*_ and its random effect ***η***_*k*_ enter the prediction only through the sum ***θ***_*k*_ +***η***_*k*_, every ***θ***_*k*_-derivative of **ŷ** equals the corresponding ***η***_*k*_ -derivative, so the mixed sensitivities in those directions collapse onto the pure random-effect ones: *d*^2^**ŷ***/d****η*** *d****θ*** onto *d*^2^**ŷ***/d****η***^2^, and *d*^3^**ŷ***/d****η***^2^*d****θ*** and *d*^3^**ŷ***/d****η*** *d****θ***^2^ onto *d*^3^**ŷ***/d****η***^3^. The same equality holds at fourth order. Under full mu-referencing of the structural parameters, the required structural prediction sensitivities reduce to pure random-effect derivatives. Structural parameters without random effects (such as *Q* and *V*_*p*_ in the case example) retain their own derivative directions; residual and covariance parameters must also be differentiated separately.

## Methods

We illustrated the method on a two-compartment oral model with first-order absorption (rate constant *KA*), fitted by the full Laplacian method to simulated pharmacokinetic data (30 subjects, six samples each over 24 h). Between-subject variability was placed on clearance, central volume, and absorption rate, while the inter-compartmental clearance *Q* and peripheral volume *V*_*p*_ were estimated as typical values only. The sparse sampling leaves the peripheral disposition weakly identified. The nine estimated parameters are the log typical values log *TV*_*CL*_, log 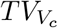, log *TV*_*KA*_, log *TV*_*Q*_, log 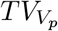, the proportional residual standard deviation *σ*, and the random-effect variances 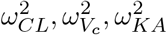. As a result, they span a wide range of curvatures, from the sharply determined *σ* to the near-flat *V*_*p*_. Structural-parameter standard errors are reported on the natural-log scale; those for *σ* and the random-effect variances are reported on their original scales.

A Python implementation of the closed-form model, using JAX [8] for sensitivities through fourth order, was used for the case example. At the fitted optimum, the analytic observed information is assembled from the saved empirical Bayes estimates using (8)–(13), without further individual optimization.

Figure 1 compares, for three parameters spanning the identifiability range, the analytic observed-information standard error of (9)–(13) with those obtained by differentiating the reconverged objective with a forward and central second difference, as the relative step *h* is swept. The comparison is repeated at two inner-mode solver settings, drawn on a common vertical scale within each row: a tight tolerance (left column; inner-mode gradient below 10^−10^) and a loose tolerance (right column; gradient below 10^−4^), at the same tightly fitted population optimum. The full set of nine parameters is shown in the Appendix, with panel (a) in Figure A1 and panel (b) in Figure A2.

**Fig 1.**
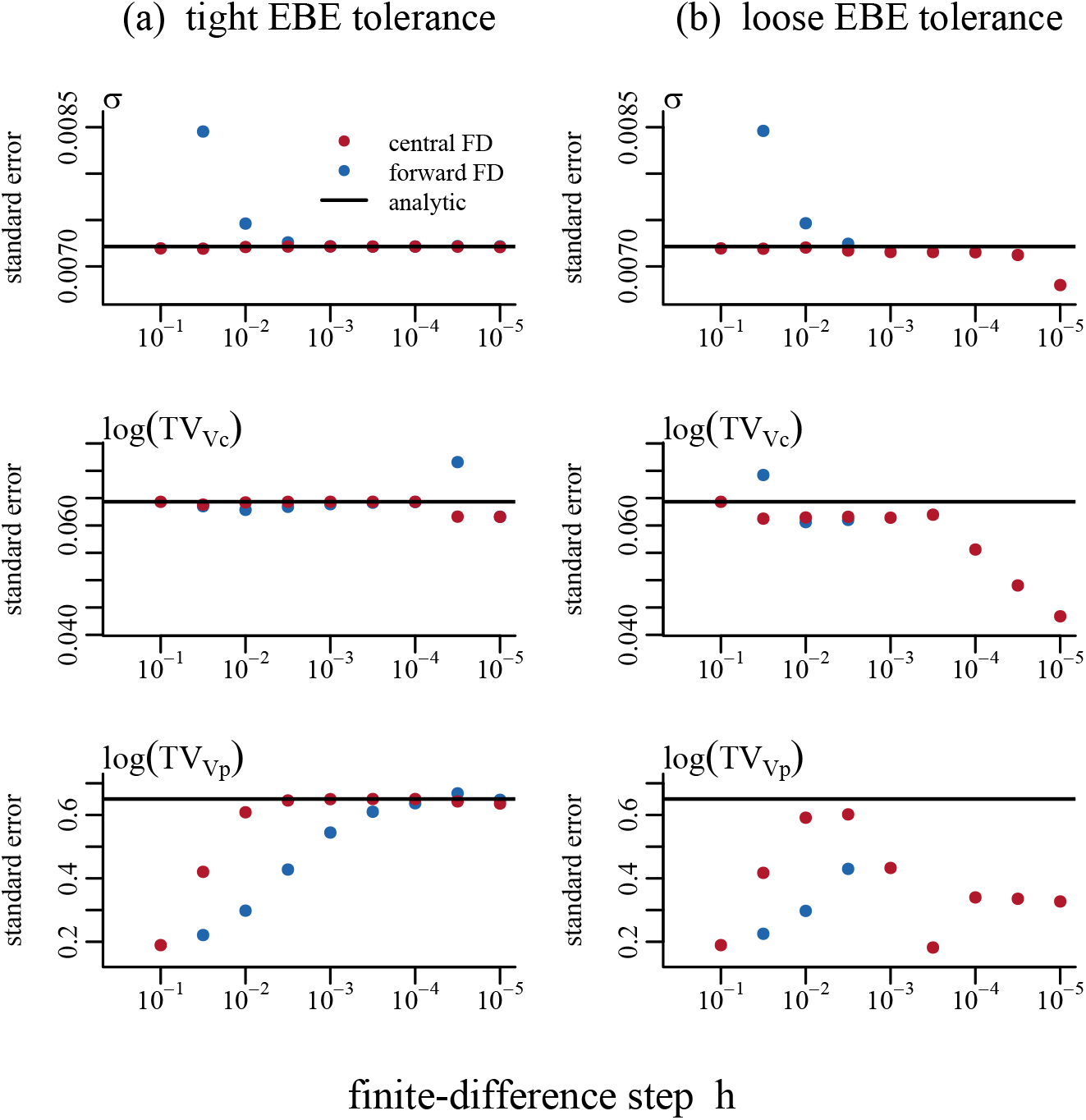
Standard errors at a common Laplacian optimum. Rows: residual SD *σ*, log 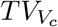 and log 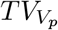. Columns: tight (10^−10^) and loose (10^−4^) EBE gradient tolerances in perturbed evaluations. Central second differences (red), forward second differences (blue), fourth-order automatic differentiation reference (black). Missing points indicate an invalid differenced information matrix. Analytic covariance uses fitted EBEs without reoptimization. All nine parameters appear in Figures A1 and A2. EBE, empirical Bayes estimate; FD, finite difference.

## Results

The analytic standard error is the reference (the solid line), while the finite differences track the line only over an intermediate band of steps whose width is parameter dependent. Loosening the inner-mode solver tolerance disturbs this comparison: at *h* = 10^−3^, the central-difference standard error for log 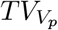 is 0.650 at the tight tolerance but 0.433 at the loose tolerance, against the analytic reference of 0.650.

The analytic covariance requires solving no perturbed empirical Bayes optimization problems. None failed in this example; avoiding them removes a possible cause of covariance-step failure after successful estimation. It does not guarantee a positive-definite population information matrix.

## Discussion

In this study we aimed to give the observed information of the Laplacian population likelihood family in closed form, obtained by continuing the exact-gradient construction by one differentiation. The observed information requires higher-order sensitivities in precisely the log-determinant term *T* ^*π*^ of (12), and the third- and fourth-order sensitivities are confined there. Under FOCE/FOCEI, third order enters through the explicit 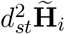 of (13) and the second mode derivative (10); under the full Laplacian method, fourth order is additionally needed for 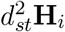. When the inner Hessian is fixed (its parameter and mode dependence ignored), *T* ^*π*^ vanishes and the population Hessian reduces to the summed data term (11), which requires nothing beyond the second-order quantities of the gradient. The covariance of this family is thus obtained by continuing the exact-gradient construction by one differentiation, the only higher-order addition being the log-determinant curvature (12)–(13) and the third- and fourth-order sensitivity equations (20)–(21) that supply it.

Two practical consequences follow. First, a procedure that already computes the exact gradient reuses its second-order machinery, so the data term (11) adds only *∂*^2^*l*_*i*_*/∂****θ***^2^; the third- and fourth-order sensitivities supply the log-determinant curvature. Higher-order sensitivities can be generated by symbolic differentiation or automatic differentiation. Forward-mode work follows input directions and reverse-mode work follows selected outputs, so their relative dimensions guide the choice of forward, reverse or mixed mode. Second, the analytic observed information is not subject to the step-dependent error of a differenced Hessian, so its standard errors are limited by the accuracy of the mode, sensitivity and differential-equation solutions and conditioning, with no step size to tune. The covariance step uses the fitted empirical Bayes estimates without reoptimization, removing failed perturbed individual optimization problems as a cause of covariance-step failure after a successful fit.

A less demanding implementation is available when the exact highest-order sensitivities are inconvenient to assemble: the required third-order sensitivities for FOCE/FOCEI can be approximated by finite-differencing the analytic second-order sensitivities in ***η*** and ***θ***, leaving the two lower orders exact. For the full Laplacian method, fourth-order sensitivities can similarly be approximated by differencing analytic third-order sensitivities, or by taking second finite differences of analytic second-order sensitivities. The latter amplifies sensitivity evaluation error more strongly as the step decreases. Neither route needs inner reconvergence — the *O*(*P* ^2^) reconverged objective evaluations for *P* population parameters are avoided, and every quantity stays anchored to the single converged mode. It reintroduces step sizes for the numerically differentiated orders.

The higher-order machinery assembled here is not tied to the observed information alone. The same third-order sensitivity equations connect the conditional-estimation family to the full Laplacian approximation. Under the first-order inner Hessian of FOCE and FOCEI the third order is required for the observed information; under the exact inner Hessian of the Laplacian approximation it is required one differentiation earlier, for the gradient — an extension identified as requiring third-order sensitivity equations. The machinery developed here supplies both orders: third order for the Laplacian gradient and fourth order for its observed information. The FOCEI objective and its gradients are also used beyond classical estimation. In nlmixr2, the machine-learning variational autoencoder (VAE) method [9] trains its encoder using the FOCEI inner likelihood and its gradient with respect to the random effects, and can optionally use the analytic outer gradient for parameters that are not mu-referenced.

## Conclusions

The observed information of the Laplacian population likelihood family can be evaluated in closed form, by continuing the exact-gradient construction by one differentiation. The only higher-order additions beyond those already required for the gradient are the curvature of the log-determinant term and the third- and fourth-order sensitivity equations that supply it, and the higher-order dependence is confined to that single term. The resulting standard errors can carry the same precision and reproducibility that exact gradients brought to the point estimates, limited by the accuracy of the mode, sensitivity and differential-equation solutions and conditioning rather than by the step-dependent error of a finite-difference Hessian. The covariance step uses the fitted empirical Bayes estimates without reoptimizing them at perturbed population parameters. This strengthens the inferences that depend on it — confidence intervals, identifiability assessment, and uncertainty propagation.

## Supporting information

analysis

## Acknowledgments

The authors thank Pyry Välitalo for his early guidance on the Laplacian likelihood family and on common practices and implementations in nonlinear mixed-effects modeling software.

## Declarations

### Code availability

The analytic FOCE and FOCEI observed information described here is implemented in the open-source R package nlmixr2 [10], available on CRAN (https://CRAN.R-project.org/package=nlmixr2) and at https://github.com/nlmixr2/nlmixr2.

The full Laplacian case example uses the custom Python implementation `analysis/laplace_ twocmt.py` (Online Resource 1), with JAX, NumPy and SciPy. It reads the simulated dataset `analysis/twocmt_oral.csv`, also in Online Resource 1, and writes estimates, information, standard errors, finite-difference comparisons and validation metrics under `analysis/laplace-results/`. The script records dependency versions and hashes of its source and input data. The Python implementation evaluates the fourth-order Laplacian construction independently of nlmixr2’s covariance routine.

## Appendix

### Mathematical derivations

#### Mode derivatives

To compute *T* ^*l*^, *T* ^*π*^, we need the mode derivatives. The first one is expressed in equation (8) and can also be written per ***θ*** coordinate:

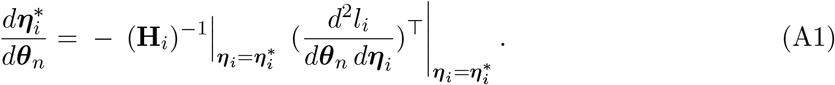

Differentiating it once more gives the second derivative,

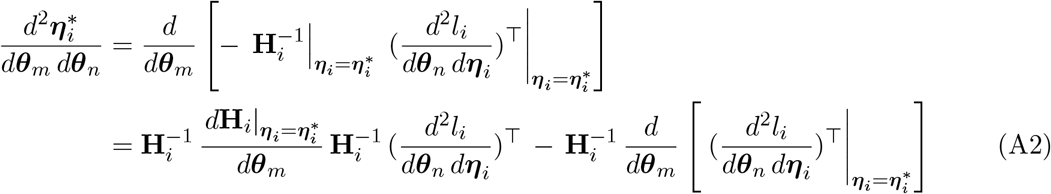

The two terms can be developed, with the first yielding

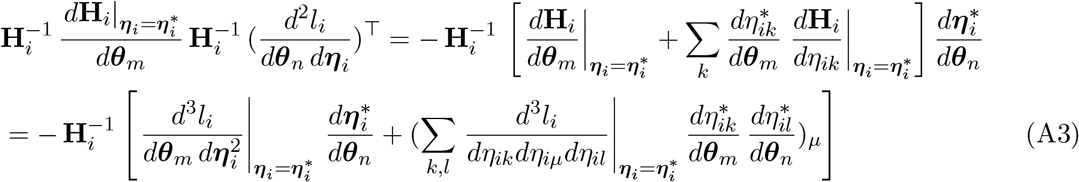

and the second yielding

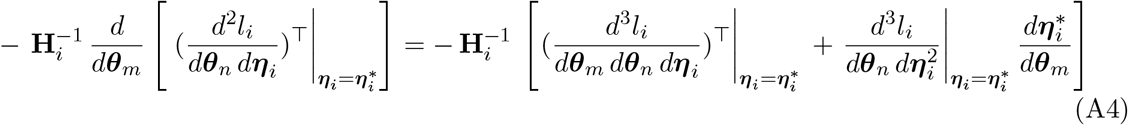

Reorganizing the terms from (A3) and (A4) recovers equation (10):

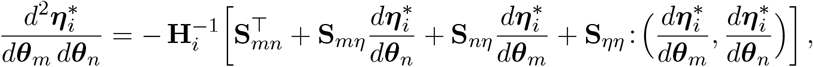

#### Data term

Since 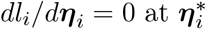, the total derivative of 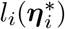 is the explicit partial *∂l*_*i*_*/∂****θ***_*m*_. Differentiating once more, we get

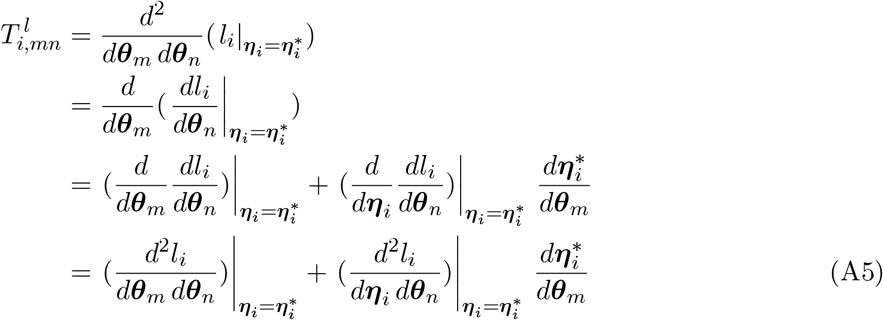

Inserting (8), we get equation (11):

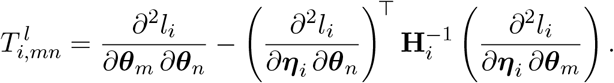

#### Log-determinant term

With *π*_*i*_ the penalty as defined in (6), 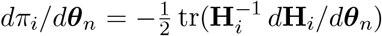 already appears in (7); where

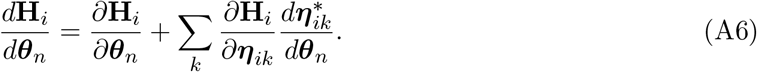

The second derivative of *π*_*i*_ is

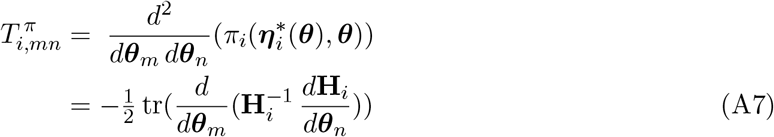

Making the total derivative explicit and using (A6), we have

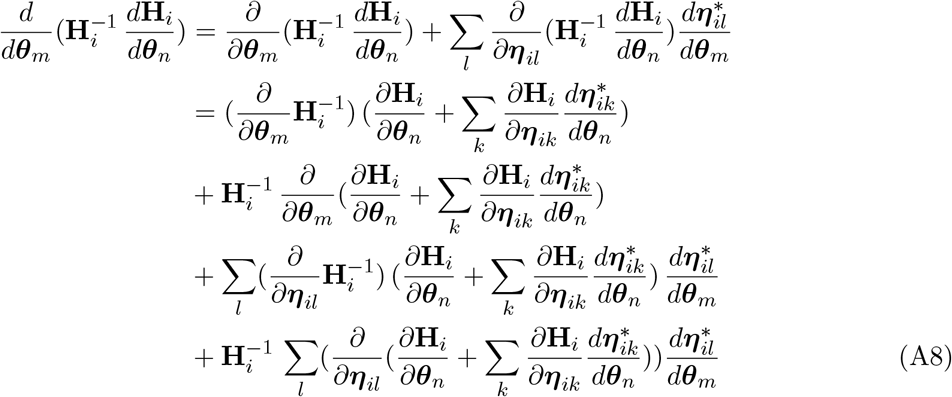

Since the mode 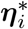 depends only on ***θ***, we have 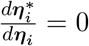 and we can group the different terms with regards to the mode derivatives

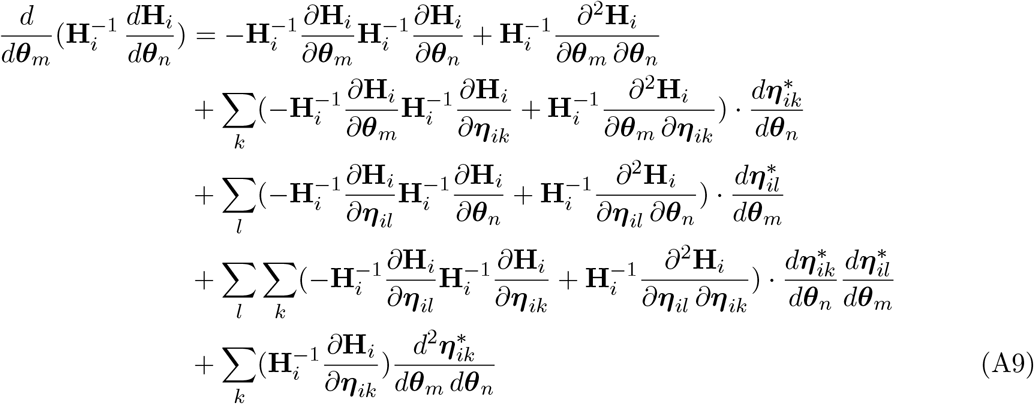

Substituting this expression into (A7) and using the linearity of the trace, we obtain equation (12):

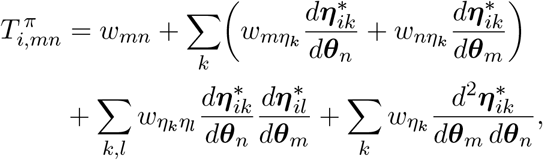

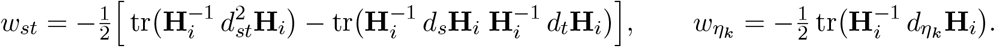

#### FOCEI Hessian expression

The Hessian in equation (15) can be rewritten by separating a prediction sensitivity term expressed in (16) and an interaction term expressed below, using the conditional second moment of ***ϵ***_*ij*_ given ***η***_*i*_.

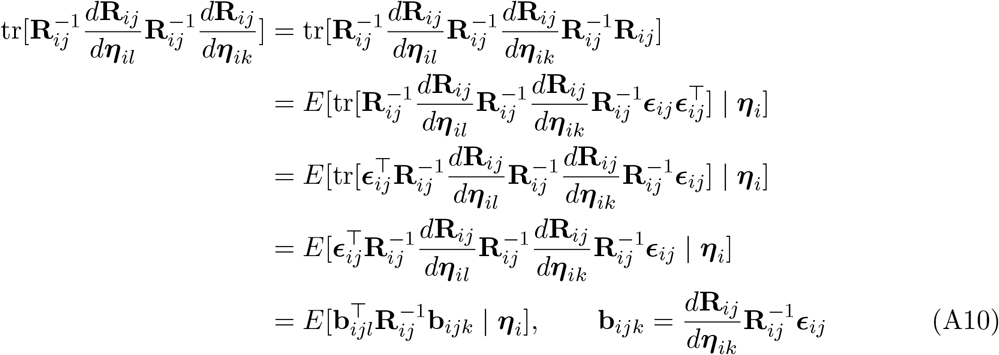

#### Further comparison with finite differences

Figures A1 and A2 repeat the comparison of Figure 1 for all nine estimated parameters, one EBE convergence tolerance per page: the tight EBE tolerance — Figure 1 panel (a) — in Figure A1, and the loose EBE tolerance — panel (b) — in Figure A2. Each parameter is drawn on a common vertical scale across the two figures, so the pages can be read against each other.

**Fig A1.**
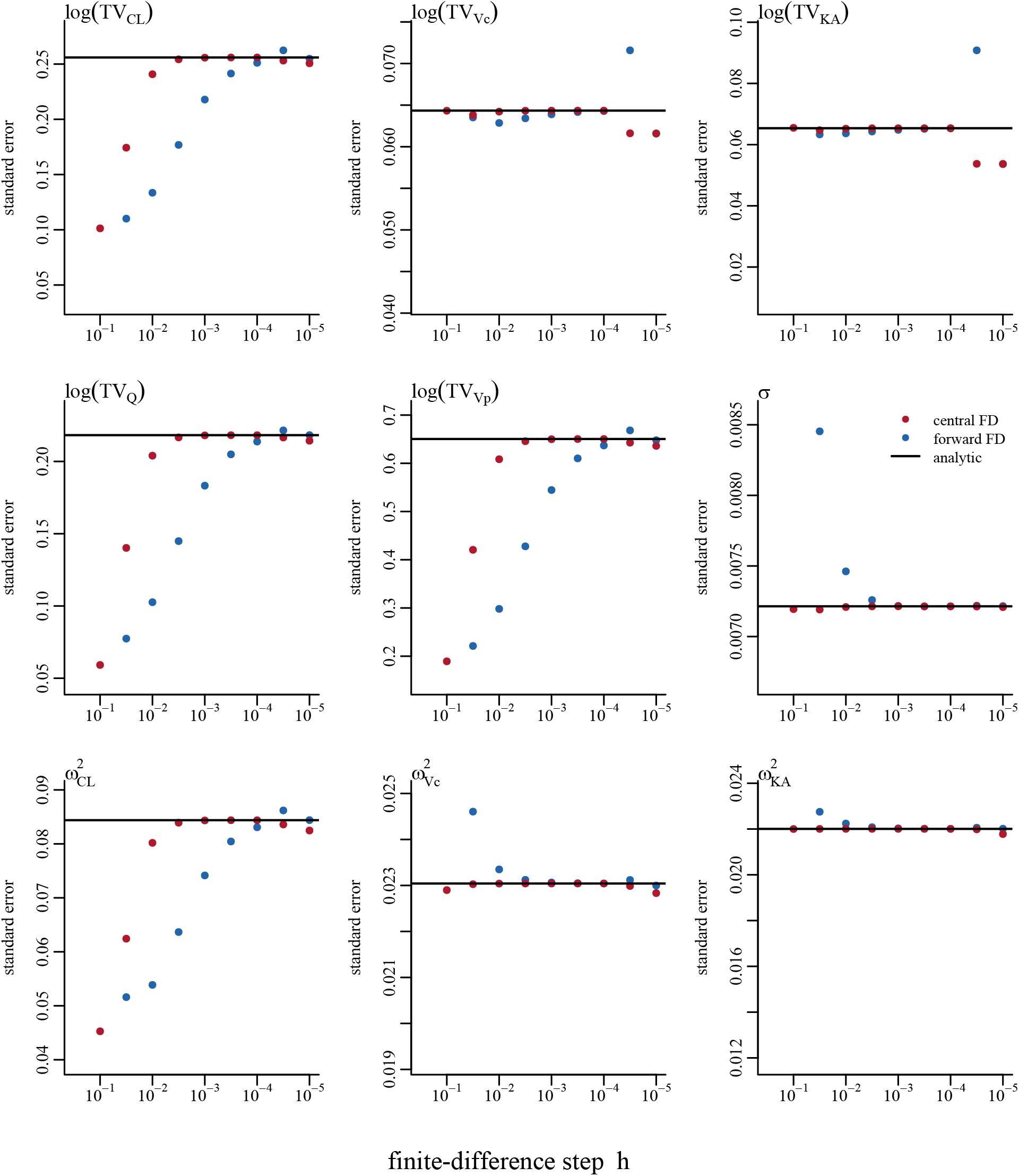
As Figure 1 panel (a), for all nine estimated parameters at tight EBE gradient tolerance (10^−10^): the log typical values log *TV*_*CL*_, log *TV*_*V*_, log *TV*_*KA*_, log *TV*_*Q*_, log 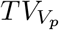, the proportional residual standard deviation *σ*, and the random-effect variances 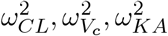. Central second difference (red), forward second difference (blue), and the fourth-order automatic differentiation Laplacian observed-information value (solid black line). EBE, empirical Bayes estimate; FD, finite difference.

**Fig A2.**
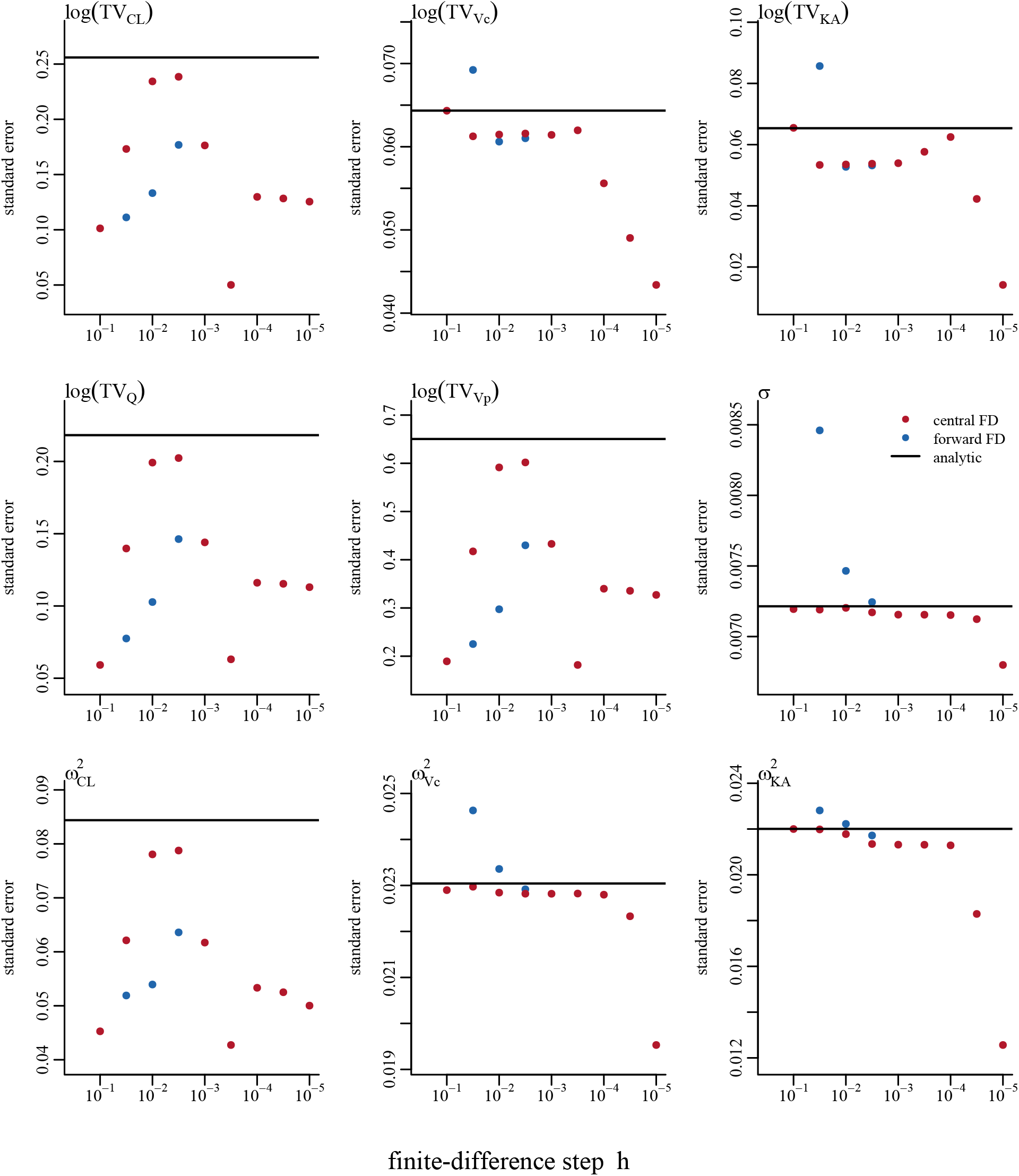
As Figure 1 panel (b), for the same nine parameters at loose EBE gradient tolerance (10^−4^), on the same per-parameter vertical scale as Figure A1. Central second difference (red), forward second difference (blue), and the fourth-order automatic differentiation Laplacian observed-information value (solid black line). EBE, empirical Bayes estimate; FD, finite difference.

